# Test-retest reliability of force plate-derived measures of reactive stepping

**DOI:** 10.1101/2020.05.10.068957

**Authors:** Tyler M. Saumur, Sunita Mathur, Jacqueline Nestico, Stephen D. Perry, George Mochizuki, Avril Mansfield

**Author notes:** Correspondence to: Tyler M. Saumur, Rehabilitation Sciences Institute, University of Toronto, 160-500 University Ave, Toronto ON M5G 1V7, Canada.

## Abstract

**Background:** Characterizing reactive stepping is important to describe the response’s effectiveness. Measures of reactive stepping related to step initiation, execution, and termination phases have been frequently reported to characterize reactive balance control. However, the test-retest reliability of these measures are unknown.

**Research questions:** What is the between- and within-session test-retest reliability of various force plate-derived measures of reactive stepping?

**Methods:** Nineteen young, healthy adults responded to 6 small (~8-10% of body weight) and 6 large perturbations (~13-15% of body weight) using an anterior lean-and-release system. Tests were conducted on two visits separated by at least two days. Participants were instructed to recover balance in as few steps as possible. Step onset, foot-off, swing, and restabilization times were extracted from force plates. Relative test-retest reliability was determined through intraclass correlation coefficients (ICCs) and 95% confidence intervals (CIs). Absolute test-retest reliability was assessed using the standard error of the measurement (SEM).

**Results:** Foot-off and swing times had the highest between- and within-session test-retest reliabilities regardless of perturbation size (between-session ICC=0.898–0.942; within-session ICC=0.455–0.753). Conversely, step onset and restabilization time had lower ICCs and wider CIs (between-session ICC=0.495–0.825; within-session ICC=−0.040–0.174). Between-session test-retest reliability was higher (ICC=0.495-0.942) for all measures than within-session test-retest reliability (ICC=−0.040–0.753). SEMs were low (3–10% of mean) for all measures, except time to restabilization (SEM=15-20% of mean), indicating good absolute reliability.

**Significance:** These findings suggest multiple baseline sessions are needed for measuring restabilization and step onset times. The SEMs provide an index for measuring meaningful change due to an intervention.

## 1. Introduction

Balance deficits are one of the most important risk factors for falls (Masud and Morris, 2001; Rubenstein, 2006). In particular, reactive balance control is required to respond to postural perturbations and necessitates rapid onset latencies to react appropriately (Mcllroy and Maki, 1995; Nashner and Cordo, 1981). When large perturbations occur, change-in-support reactions, like taking reactive steps, are the only strategy that can lead to successful stabilization (Maki and Mcilroy, 1997). Training reactive stepping has been shown to decrease the incidence of falls (Mansfield et al., 2015a). While clinical tools are available to assess reactive balance control such as the Balance Evaluation Systems Test (BESTest) (Horak et al., 2008) or the Performance-Oriented Mobility Assessment (POMA) (Tinetti, 1986), concerns exist related to the standardization of perturbation size and scoring of balance performance (Inness, 2015).

Temporally and spatially sensitive features of reactive control can affect responses and have been shown to be associated with falls (Brauer et al., 2001; Mansfield et al., 2015b). Therefore, quantitative assessments of reactive balance control may be useful for the accurate and reliable assessment of reactive stepping effectiveness.

The lean-and-release test is a method that standardizes the size and direction of a perturbation. The lean-and-release test can simulate a fall in any direction; however, the forward direction is most commonly used (Hsiao-Wecksler, 2008). The method involves an individual being connected to a cable on a wall behind them and leaning forward (Figure 1). The cable is then released once a specific amount of body weight is supported by it, resulting in a stepping response. While balance can be scored based on the success of the response (e.g., single step, multi-step, assisted step, no step (Inness et al., 2015)), additional quantitative measures can be obtained to understand the quality of the stepping response. Some of the more common measures studied during reactive stepping following forward perturbations include step onset, foot-off time, swing time (Lakhani et al., 2011; Mansfield et al., 2013; Singer et al., 2019; Tseng et al., 2009), and time to restabilization (Singer et al., 2019). These measures represent the different phases involved in a reactive step: step initiation and limb unloading, swing phase, and foot contact and restabilization (Mcllroy and Maki, 1995); furthermore, they can be obtained from force plates, which offer a simpler setup and can be more cost-effective than kinematic measures. While previous work has established test-retest reliability of force plate-derived measures during voluntary stepping (Melzer et al., 2007), these measures have not been assessed in the context of reactive stepping. Reliability of these measures is particularly important when using reactive stepping outcomes for balance training interventions. Therefore, investigating the test-retest reliability of these force plate-derived measures during reactive stepping can provide information regarding their acceptability for longitudinal testing.

**Figure 1.**
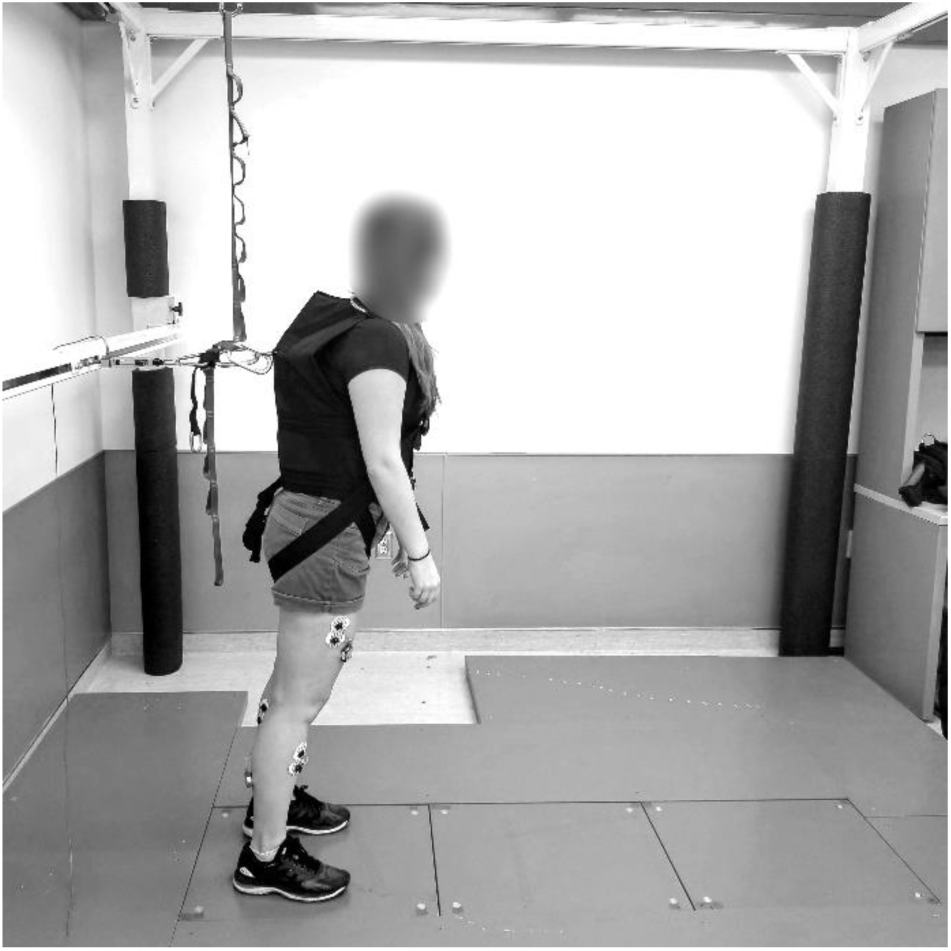
Example of experimental setup.

Accordingly, the objective of this study was to determine the between- and within-session test-retest reliability of force plate-derived measures of reactive stepping throughout the different phases of reactive stepping. We hypothesized that these measures would demonstrate moderate-to-good reliability for large and small perturbation sizes (ICC: 0.5-0.9) (Koo and Li, 2016) aligning with previous centre of pressure (COP) measures observed during voluntary stepping (Melzer et al., 2007).

## 2. Methods

### 2.1. Participants

Participants were included if they: were between the ages of 20 and 35 years old, self-reported as having no walking difficulties, were able to understand English, had no neurological or musculoskeletal disorders, had normal or corrected to normal vision, and had no health conditions that limit the ability to complete daily activities independently. Following a sample size calculation using an expected ICC of 0.6 (Melzer et al., 2007), 80% power, and an alpha of 0.05, we determined that 15-20 participants were needed (Bujang and Baharum, 2017). All participants were recruited from the community and provided written consent. Research ethics approval was obtained from Sunnybrook Research Institute (#017-2015) and the University of Toronto (#31805).

### 2.2. Instrumentation and study procedures

A lean-and-release system was used to study reactive stepping (Inness et al., 2015). Participants stood on a single force plate (referred to as the ‘standing’ plate, Type 9260AA, Kistler, Winterthur, Switzerland) in a pre-defined stance with heel centres 17 cm apart and externally rotated 14° (McIlroy and Maki, 1997). Participants were placed in a harness and leaned forward at the ankle joint with ~8-10% or ~13-15% of their body weight supported by a cable connected to the rear wall behind them (Inness et al., 2015) measured by an in-line load cell (Transducer Techniques, Temecula, USA). These perturbation magnitudes were used as they were tolerated well by participants during pilot testing, while also resulting in different response characteristics (e.g., longer steps and increased braking forces required for large perturbation magnitude). Body weight percentage was monitored in real-time and adjusted by increasing or decreasing the rear cable length for each trial accordingly. For each trial, once the participant was leaning into the harness and stated that they were ready, the investigator released the cable at a random time-point 1 to 10 s later, resulting in participants having to take a reactive step onto the force plate(s) (referred to as the ‘stepping plate’) to regain balance. Participants were instructed to “recover their balance in as few steps as possible” with the specified leg. At the start of each session, participants performed a single, initial small perturbation where they were not given any information regarding which foot to step with. The foot used for this initial response was characterized as the preferred stepping limb. This initial trial was not included in the analysis. Following the initial perturbation trial, three trials were performed for each foot and condition (small/large perturbation) resulting in 12 trials in total. Participants performed one block of perturbations per stepping limb (one block for preferred limb, one block for non-preferred limb). Perturbation size was randomized within each block of trials. Participants completed two lab sessions separated by at least two days and no more than ten days (Figure 2). Randomization was achieved using the random function in Microsoft Excel. Medial/lateral (ML) COP position was monitored online to ensure anticipatory offloading of the stepping limb did not occur prior to perturbation onset. Participants were instructed to relax and centre their weight if anticipatory offloading was evident. Following each perturbation, participants were asked to hold their position for 7 s.

**Figure 2.**
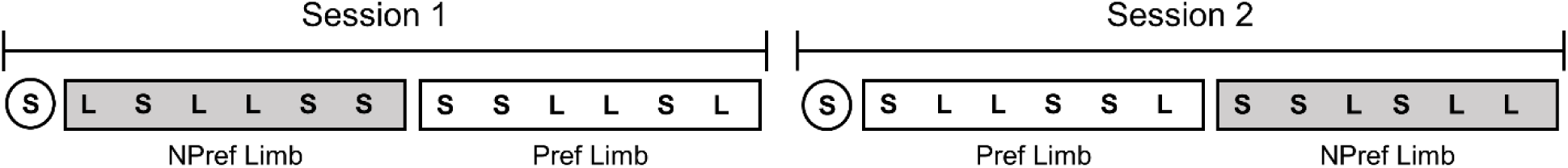
Example of a participant’s two visits. An initial small perturbation was performed to determine the individual’s preferred stepping limb. Two blocks of trials separated based on limb were then completed, with perturbation size randomized. In the second session, an initial small perturbation was presented again, followed by two blocks counterbalanced for limb order. S, small perturbation; L, large perturbation

### 2.3. Questionnaires

During their first visit, participants completed three questionnaires to gauge their balance confidence, physical activity levels, and lower limb dominance. Participants completed the Activities-Specific Balance Confidence (ABC) Scale, a 16-item questionnaire, where they rate their confidence in maintaining balance during various situations on a scale from 0-100% (Powell and Myers, 1995). Participants also filled out the International Physical Activity Questionnaire (IPAQ) short form, which is a one-page questionnaire that characterizes individuals’ involvement in physical activity over the previous seven days (Craig et al., 2003). In addition, participants completed the Waterloo Footedness Questionnaire to characterize their preferred lower limb for various tasks (Elias et al., 1998). For the present study, questions 2, 4, 6, 8, and 10 were used to determine the preferred support limb as these questions pertain to balance tasks.

### 2.4. Data processing

Data were acquired using a Power 1401 mkII (Cambridge Electronic Design, Cambridge, UK). All force data were collected using data acquisition software (Spike2 Version 7.17, Cambridge Electronic Design, Cambridge, UK). Ground reaction forces and load cell data were sampled at 256 Hz and stored for offline analysis. COP data were low-pass filtered offline using a fourth order dual pass Butterworth filter at 10 Hz. ML-COP position was calculated for the standing plate for force plate measures prior to foot contact. Net anterior/posterior (AP)-COP was calculated from each force plate and the net AP-COP was then derived to calculate AP-COP velocity for restabilization measures.

### 2.5. Measures

Figure 3A outlines the measures prior to foot contact. Time to step onset was defined as the time from perturbation onset (< 5 N on load cell) to the point when the ML-COP deviated more than 4 mm from the mean ML-COP calculated over 500 ms prior to perturbation onset (McIlroy and Maki, 1999). As anticipatory postural adjustments (APAs) can affect step onset time (MacKinnon et al., 2007), they were also characterized. Foot-off time was defined as the time from perturbation onset to when the ML-COP was 20% of the max slope when shifting towards the stepping limb (Kurz et al., 2013). Swing time occurred from foot-off to foot contact (>5 N on stepping plate). Time to restabilization was calculated as the time from foot contact to the time at which the net AP-COP velocity returned to within 3 standard deviations of the mean net AP-COP velocity during the last 2 s of foot contact for at least 1 s (Figure 3B). All data were visually inspected and corrected manually when appropriate.

**Figure 3.**
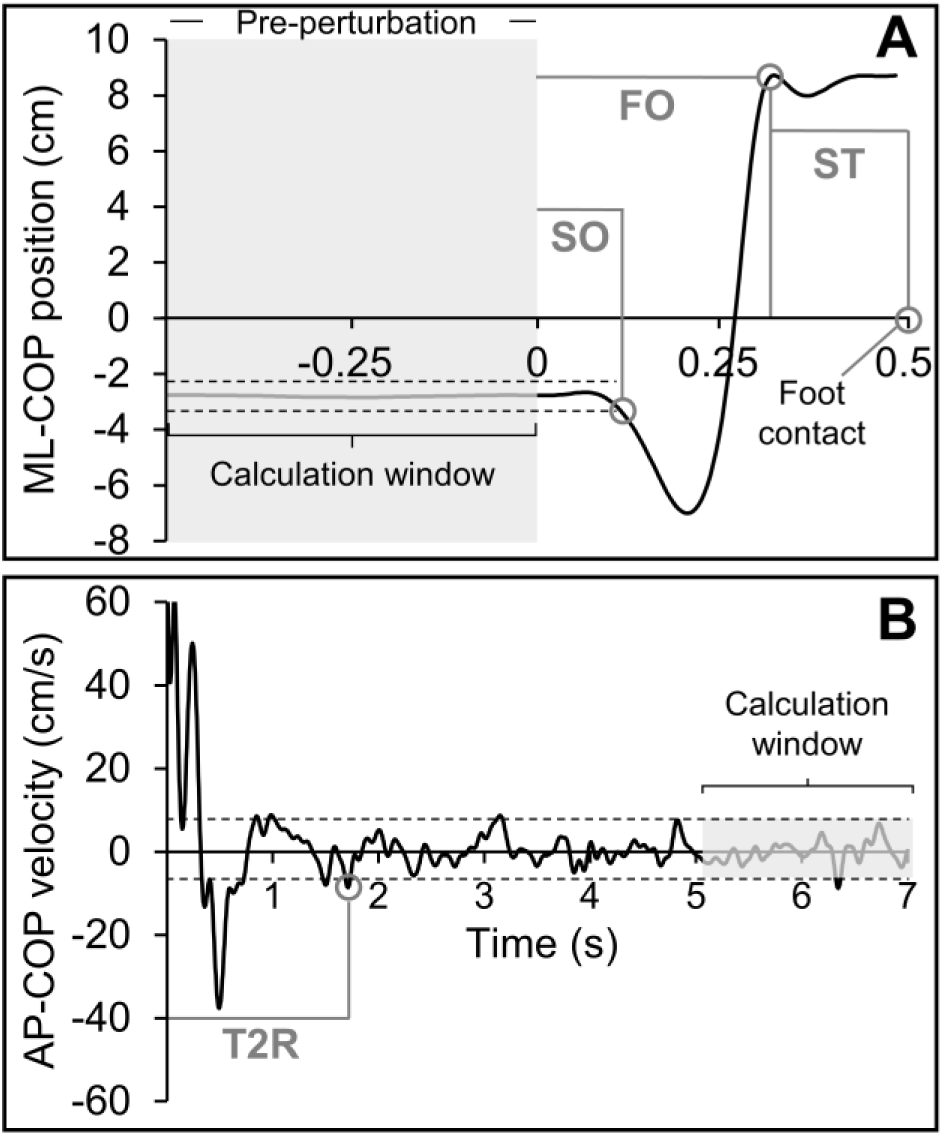
Representative reactive step response for one participant. **A.** Force plate measures taken prior to foot contact from ML-COP position. Time 0 is perturbation onset. **B.** AP-COP velocity following foot contact. Restabilization is achieved when the AP-COP velocity enters the bandwidth and stays within it for at least 1 s. Time starts at foot contact. SO, step onset; FO, foot-off; ST, swing time; T2R, time to restabilization.

### 2.6. Statistical analysis

To determine the test-retest reliability, ICCs between the two sessions were then calculated using two-way mixed effects (2,1) models with absolute agreement using the average measures of all trials for each perturbation size to provide a relative measure of reliability (Weir, 2005). The absolute reliability for each measure between sessions was also evaluated by calculating the standard error of the measurement (SEM) (Weir, 2005). The SEM was estimated as √*Ms*_*E*_ from a two-way ANOVA (Hopkins, 2000). This method provides an absolute measure of reliability, while avoiding errors associated with calculating SEM from the ICC (Weir, 2005). To visualize the agreement between the two sessions, Bland–Altman plots were created for all measures. To test within-session reliability, ICCs were determined for each session using a two-way mixed effects (2,1) model with absolute agreement using the single measures from all trials for each perturbation size, with all trials being compared to each other. Reliability was categorized as poor (<0.5), moderate (0.5 to 0.75), good (>0.75 to 0.9), or excellent (>0.9) based on the ICCs and 95% CIs (Koo and Li, 2016). Values are reported as mean (standard error of the mean) unless otherwise stated. Significance was set to *p* < 0.05.

## 3. Results

Twenty young, healthy adults participated in this study. Data from one participant was excluded from analysis as they failed to reach restabilization on more than half the trials, therefore 19 participants were included in the analysis; see Table 1 for characteristics of these participants. The mean length of time between sessions was 5.8 days (standard deviation: 2.5). A total of 450 trials were analyzed, with 6 being removed due to excessive movement following the reactive step or stability unable to be obtained on the force plates. The mean (standard error of the mean) of large perturbation sizes for session 1 and session 2 were 13.7 (0.2)% and 13.9 (0.3)% of body mass, respectively. Mean small perturbation sizes for session 1 and 2 were 10.1 (0.2)% and 9.9 (0.2)% of body mass, respectively. The number (proportion) of trials where APAs were present are observed in Table 2.

**Table 1.**
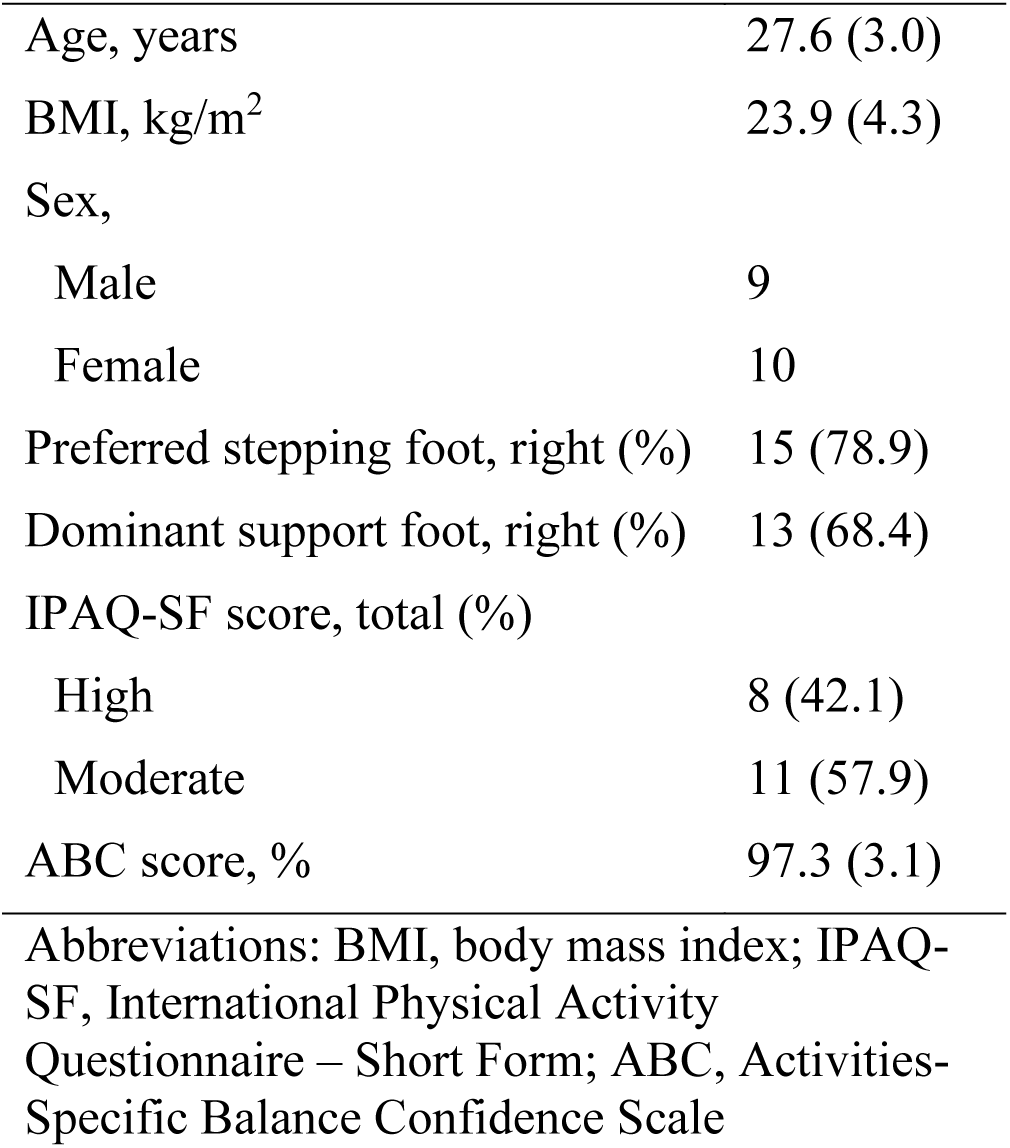
Study participant characteristics (n = 19). Values presented are means with standard deviations in parentheses (continuous variables) or counts.

|  |  |
| --- | --- |
| Age, years | 27.6 (3.0) |
| BMI, kg/m<sup>2</sup> | 23.9 (4.3) |
| Sex, |  |
| Male | 9 |
| Female | 10 |
| Preferred stepping foot, right (%) | 15 (78.9) |
| Dominant support foot, right (%) | 13 (68.4) |
| IPAQ-SF score, total (%) |  |
| High | 8 (42.1) |
| Moderate | 11 (57.9) |
| ABC score, % | 97.3 (3.1) |

Abbreviations: BMI, body mass index; IPAQ-SF, International Physical Activity Questionnaire – Short Form; ABC, Activities-Specific Balance Confidence Scale

**Table 2.** Presence of APAs in trials. Values shown as count (% of total). Both large and small perturbation trials had 111 and 114 trials analyzed for session 1 and 2, respectively.

|  | Large perturbations |  |  | Small perturbations |  |  |
| --- | --- | --- | --- | --- | --- | --- |
|  | Single APA | Multi APA | No APA | Single APA | Multi APA | No APA |
| Session 1 | 100 (90.1%) | 4 (3.6%) | 7 (6.3%) | 106 (95.5%) | 1 (0.9%) | 4 (3.6%) |
| Session 2 | 100 (87.7%) | 3 (2.6%) | 11 (9.7%) | 107 (93.9%) | 4 (3.5%) | 3 (2.6%) |

Time to step onset (small perturbations only), foot-off and swing time demonstrated good to excellent test-retest reliability between sessions (ICC = 0.825-0.942; Table 3) when taking into account the 95% CIs (Koo and Li, 2016). Conversely, time to step onset (large perturbations) and time to restabilization (for both large and small perturbations) exhibited wide CIs ranging from poor to good between-session reliability. The SEMs for all measures are reported in Table 3. In general, SEMs were small relative to the means (3-10%), except for time to restabilization which demonstrated SEMs that were approximately 15-20% of the mean. Upon inspection of the Bland-Altman plots (Figure 4), all biases are approximately 0 (range: −0.054–0.233) with consistent variability as means increase. Furthermore, perturbation magnitude appeared to affect the limits of agreement for step onset and time to restabilization, which had wider limits of agreements during large perturbation trials. Fourteen participants with full data sets were included in the within-session reliability analysis. Measures were not consistently reliable based on perturbation size and session (Table 4). In general, foot-off and swing time demonstrated the best within-session reliability with ICCs ranging from 0.455 to 0.753, however 95% CIs were wide.

**Table 3.** Between-session reliability of the force plate-derived measures. Values for sessions 1 and 2 are reported as mean (standard error of the mean) in seconds (s).

|  | Session 1 (s) | Session 2 (s) | SEM (s) | ICC<sub>[2,1]</sub> [95% CI] |
| --- | --- | --- | --- | --- |
| Large perturbation |  |  |  |  |
| SO | 0.113 (0.003) | 0.115 (0.003) | 0.011 | 0.548 [-0.198, 0.827] |
| FO | 0.309 (0.007) | 0.306 (0.006) | 0.010 | 0.933 [0.828, 0.974] |
| ST | 0.176 (0.006) | 0.168 (0.007) | 0.011 | 0.898 [0.700, 0.962] |
| T2R | 2.139 (0.129) | 1.906 (0.112) | 0.424 | 0.495 [-0.205, 0.798] |
| Small perturbation |  |  |  |  |
| SO | 0.119 (0.003) | 0.116 (0.003) | 0.006 | 0.825 [0.558, 0.932] |
| FO | 0.350 (0.010) | 0.349 (0.010) | 0.015 | 0.942 [0.848, 0.978] |
| ST | 0.179 (0.009) | 0.169 (0.008) | 0.013 | 0.920 [0.760, 0.971] |
| T2R | 1.739 (0.102) | 1.793 (0.097) | 0.281 | 0.741 [0.323, 0.900] |

SO, step onset; FO, foot-off; ST, swing time; T2R, time to restabilization; SEM, standard error of measurement

**Table 4.**
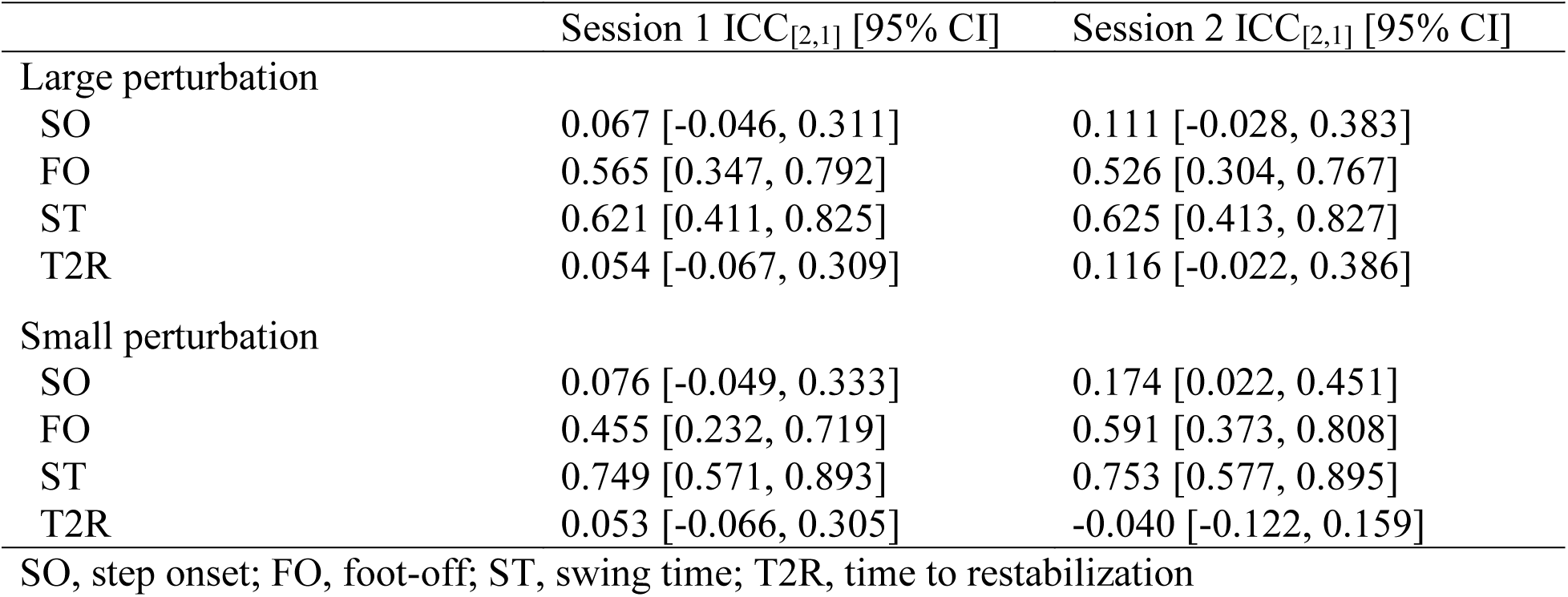
Within-session reliability of force plate-derived measures for sessions 1 and 2

**Figure 4.**
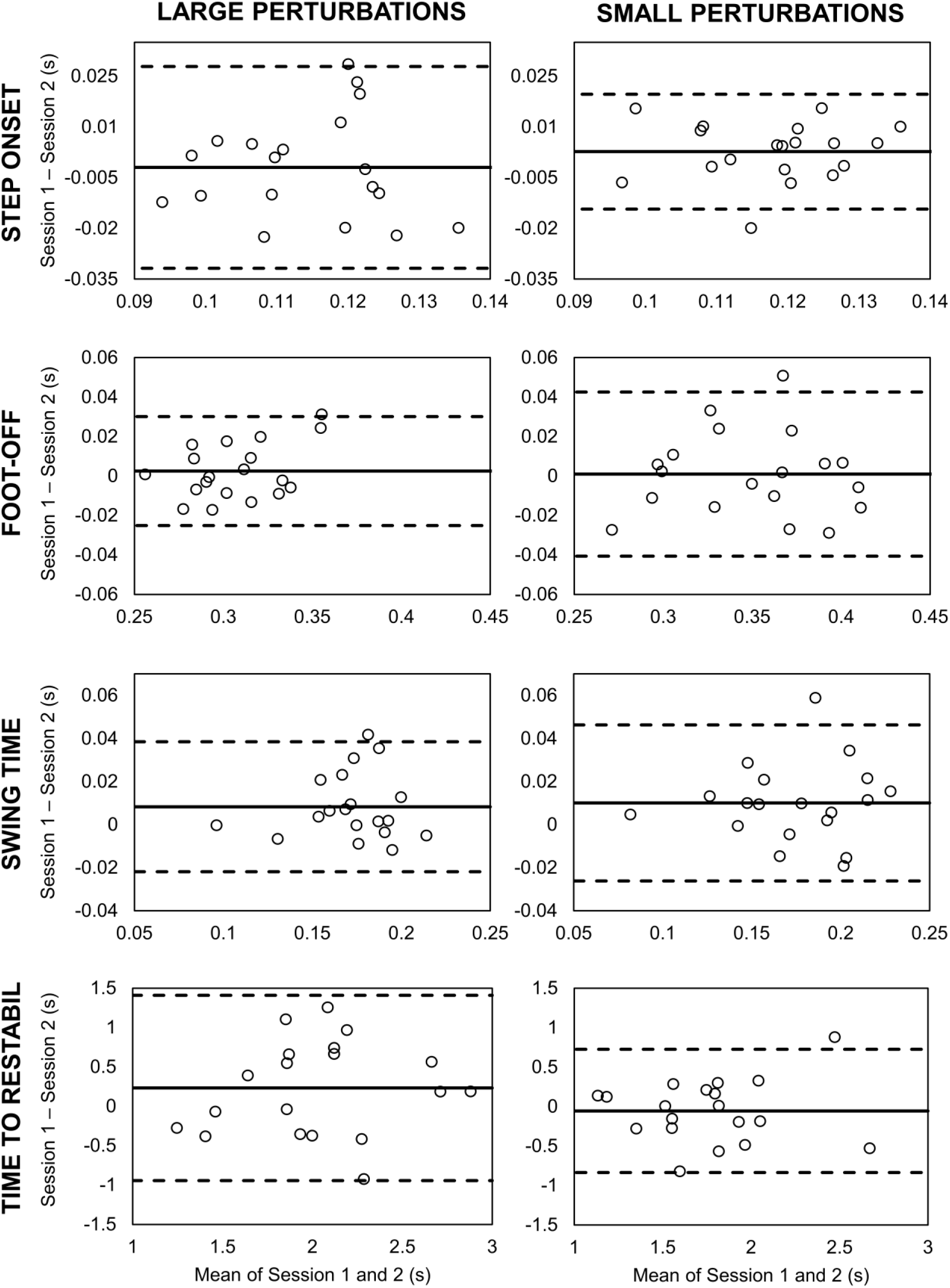
Bland-Altman plots for all measures during both perturbation magnitudes.

## 4. Discussion

The purpose of this study was to determine the test-retest reliability of force plate-derived measures during reactive stepping. We found that foot-off and swing time demonstrated the best between- and within-session test-retest reliability regardless of perturbation size and that time to step onset and time to restabilization exhibited large CIs ranging from poor to excellent test-retest reliability. SEMs were low (3-10% of mean) for all measures except for time to restabilization, indicating good relative reliability.

The measures that demonstrated the strongest between- and within-session test-retest reliability irrespective of condition (large or small perturbation) were foot-off time and swing time. Interestingly, previous work in voluntary stepping has found foot-off time to have excellent reliability, but not swing time (Melzer et al., 2007). This may be due to the limited time available for the stepping leg to rapidly catch the centre of mass (COM) following a forward perturbation, resulting in more similar values between sessions and tighter CIs. Indeed, Lakhani and colleagues measured step onset, foot-off time, and swing time during lean-and-release perturbations and swing time was the measure that demonstrated the smallest coefficient of variation within a single session (Lakhani et al., 2011). Our values also appear to be fairly similar to those previously reported (Lakhani et al., 2011; Singer et al., 2019). In the present study, swing time also appeared to be relatively consistent regardless of perturbation size, whereas foot-off time may have to account for the larger perturbation magnitude as it was faster during these trials. Perturbation size appeared to have no impact on the reliability for either measure.

While foot-off and swing times demonstrated good to excellent test-retest reliability, this was not evident for time to step onset. This is in contrast with previous literature in voluntary stepping, whereby good reliability has been reported for step onset time (Melzer et al., 2007). Interestingly, time to step onset demonstrated moderate to excellent reliability for the small, but not the large perturbations. This is further supported by wider limits of agreement during the large perturbation trials, suggesting greater variability between sessions. One potential reason for this could be a greater number of multi APA and no APA trials in the large compared to the small perturbation trials. This variability in anticipatory control for the large perturbation condition may provide a potential explanation for this discrepancy, as APAs impact step onset time (MacKinnon et al., 2007). Time to restabilization also demonstrated poor test-retest reliability which may be due to the variability that can occur in individuals’ behavioural responses to balance perturbations. While the timing of step initiation and execution are locked, the amount of time needed to restabilize may be more variable. Furthermore, as time to restabilization occurs after the step, how the step is initiated and executed will impact the length of time needed to restabilize. This is supported by work showing that shorter step length and higher AP-COM velocity is characteristic of multi-step responses, whereby restabilization does not occur on the initial reactive step (McIlroy and Maki, 1996). Interestingly, our time to restabilization values were faster than those previously observed (Singer et al., 2019). The predictable nature of our protocol may have allowed for increased preparation time, potentially leading to faster restabilization times. It is worth noting that time to restabilization in reactive stepping has been previously studied kinematically using AP-COM velocity (Singer et al., 2019). To date, there is no evidence suggesting that these differences could be related to different control mechanisms of the COP and COM velocity. Masani et al. (2014) have found that COP velocity is more strongly correlated with COM acceleration than velocity during quiet standing [9]; however, this relationship has not been explored during reactive stepping. Future work should explore the relationship between COM and COP velocity following reactive stepping to understand the associations between these biomechanical variables.

This work is not without its limitations. As this work was conducted on young, healthy individuals, the reliability of the measures may not translate to older adults or those with balance impairments. Both the direction and size of the perturbations were predictable, as all perturbations resulted in a forward step and perturbation magnitude could be anticipated based on the degree of lean. However, we randomized the perturbation size to minimize any potential for habituation. To further understand the reliability of force plate-derived measures of reactive balance control, both blocked and randomized perturbation designs should be conducted for large and small perturbation magnitudes to understand how habituation affects reliability. Future work should also compare the restabilization of AP-COP velocity to AP-COM velocity to further elucidate the relationship between the two measures.

In conclusion, this study found that foot-off and swing time have good to excellent test-retest reliability based on ICCs, and all measures have good absolute reliability, except for time to restabilization. Based on these findings, multiple baseline sessions are recommended if measuring time to restabilization and potentially step onset changes over time. These SEMs will also provide an index for measuring meaningful change within individuals that may occur due to aging or an intervention. Studying the reliability of these measures in different clinical populations and older adults will further inform our ability to use these variables to assess changes in reactive balance control longitudinally.

## ACKNOWLEDGEMENTS

The investigators would like to thank all participants for their dedication and time. TS was supported by the Ontario Graduate Scholarship, Toronto Rehabilitation Institute Student Scholarship, and Peterborough KM Hunter Charitable Foundation Graduate Award. AM holds a New Investigator Award from the Canadian Institutes of Health Research (MSH-141983).

## References

Brauer, S.G., Woollacott, M., Shumway-Cook, A., 2001. Interacting effects of cognitive demand and recovery of postural stability in balance-impaired elderly persons. Journals Gerontol. Ser. A Biol. Sci. Med. Sci. 56, M489–M496. doi:10.1093/gerona/56.8.M489

Bujang, M.A., Baharum, N., 2017. A simplified guide to determination of sample size requirements for estimating the value of intraclass correlation coefficient: a review. Arch. Orofac. Sci. 12, 1–11.

Craig, C.L., Marshall, A.L., Sjöström, M., Bauman, A.E., Booth, M.L., Ainsworth, B.E., Pratt, M., Ekelund, U., Yngve, A., Sallis, J.F., Oja, P., 2003. International physical activity questionnaire: 12-Country reliability and validity. Med. Sci. Sports Exerc. 35, 1381–1395. doi:10.1249/01.MSS.0000078924.61453.FB

Elias, L.J., Bryden, M.P., Bulman-Fleming, M.B., 1998. Footedness is a better predictor than is handedness of emotional lateralization. Neuropsychologia 36, 37–43. doi:10.1016/S0028-3932(97)00107-3

Hopkins, W.G., 2000. Measures of reliability in sports medicine and science. Sport. Med. 30, 1–15.

Horak, F.B., Wrisley, D., Frank, J., 2008. The Balance Evaluations Systems Test (BESTest) to differentiate balance deficits. Phys. Ther. 89, 484–498. doi:http://dx.doi.org/10.1111/j.1467-9280.2007.01910.x

Hsiao-Wecksler, E.T., 2008. Biomechanical and age-related differences in balance recovery using the tether-release method. J. Electromyogr. Kinesiol. 18, 179–187. doi:10.1016/j.jelekin.2007.06.007

Inness, E.L., 2015. Reactive balance control after stroke: Towards enhanced clinical understanding and assessment. Dr. Diss.

Inness, E.L., Mansfield, A., Biasin, L., Brunton, K., Bayley, M., McIlroy, W.E., 2015. Clinical implementation of a reactive balance control assessment in a sub-acute stroke patient population using a “lean-and-release” methodology. Gait Posture 41, 529–534. doi:10.1016/j.gaitpost.2014.12.005

Koo, T.K., Li, M.Y., 2016. A guideline of selecting and reporting intraclass correlation coefficients for reliability research. J. Chiropr. Med. 15, 155–163. doi:10.1016/j.jcm.2016.02.012

Kurz, I., Berezowski, E., Melzer, I., 2013. Frontal plane instability following rapid voluntary stepping: effects of age and a concurrent cognitive task. Journals Gerontol. – Ser. A Biol. Sci. Med. Sci. 68, 1402–1408. doi:10.1093/gerona/glt040

Lakhani, B., Mansfield, A., Inness, E.L., McIlroy, W.E., 2011. Characterizing the determinants of limb preference for compensatory stepping in healthy young adults. Gait Posture 33, 200–204. doi:10.1016/j.gaitpost.2010.11.005

MacKinnon, C.D., Bissig, D., Chiusano, J., Miller, E., Rudnick, L., Jager, C., Zhang, Y., Mille, M.L., Rogers, M.W., 2007. Preparation of anticipatory postural adjustments prior to stepping. J. Neurophysiol. 97, 4368–4379. doi:10.1152/jn.01136.2006

Maki, B.E., Mcilroy, W.E., 1997. The role of limb movements in maintaining upright stance: the “change-in-support” strategy. Phys. Ther. 77, 488–507.

Mansfield, A., Inness, E.L., Wong, J.S., Fraser, J.E., Mcilroy, W.E., 2013. Is impaired control of reactive stepping related to falls during inpatient stroke rehabilitation ? Neurorehabil. Neural Repair 27, 526–533. doi:10.1177/1545968313478486

Mansfield, A., Wong, J.S., Bryce, J., Knorr, S., Patterson, K.K., 2015a. Does perturbation-based balance training prevent falls? Systematic review and meta-analysis of preliminary randomized controlled trials. Phys. Ther. 95, 700–709. doi:10.2522/ptj.20140090

Mansfield, A., Wong, J.S., McIlroy, W.E., Biasin, L., Brunton, K., Bayley, M., Inness, E.L., 2015b. Do measures of reactive balance control predict falls in people with stroke returning to the community? Physiotherapy 101, 373–380. doi:10.1016/j.physio.2015.01.009

Masud, T., Morris, R., 2001. Epidemiology of falls. Age Ageing 30, 3–7. doi:10.1093/ageing/30.suppl

McIlroy, W.E., Maki, B.E., 1999. The control of lateral stability during rapid stepping reactions evoked by antero-posterior perturbation: Does anticipatory control play a role? Gait Posture 9, 190–198. doi:10.1016/S0966-6362(99)00013-2

McIlroy, W.E., Maki, B.E., 1997. Preferred placement of the feet during quiet stance: Development of a standardized foot placement for balance testing. Clin. Biomech. 12, 66–70. doi:10.1016/S0268-0033(96)00040-X

McIlroy, W.E., Maki, B.E., 1996. Age-related changes in compensatory stepping in response to unpredictable perturbations. Journals Gerontol. – Ser. A Biol. Sci. Med. Sci. 51, 289–296. doi:10.1093/gerona/51A.6.M289

Mcllroy, W.E., Maki, B., 1995. Adaptive changes to compensatory stepping responses. Gait Posture 3, 43–50.

Melzer, I., Shtilman, I., Rosenblatt, N., Oddsson, L.I.E., 2007. Reliability of voluntary step execution behavior under single and dual task conditions. J. Neuroeng. Rehabil. 4. doi:10.1186/1743-0003-4-16

Nashner, L.M., Cordo, P., 1981. Relation of automatic postural responses and reaction-time voluntary movements of human leg muscles. Exp. Brain Res. 43, 395–405.

Powell, L.E., Myers, A.M., 1995. The Activities-specific Balance Confidence (ABC) Scale. Journals Gerontol. Ser. A Biol. Sci. Med. Sci. 50, M28–M34.

Rubenstein, L.Z., 2006. Falls in older people: epidemiology, risk factors and strategies for prevention. Age Ageing 35, 37–41. doi:10.1093/ageing/afl084

Singer, J.C., Prentice, S.D., McIlroy, W.E., 2019. Exploring the role of applied force eccentricity after foot-contact in managing anterior instability among older adults during compensatory stepping responses. Gait Posture 73, 161–167. doi:10.1016/j.gaitpost.2019.07.250

Tinetti, M.E., 1986. Performance-oriented assessment of mobility problems in elderly patients. J. Am. Geriatr. Soc. 34, 119–126. doi:10.1111/j.1532-5415.1986.tb05480.x

Tseng, S.C., Stanhope, S.J., Morton, S.M., 2009. Impaired reactive stepping adjustments in older adults. Journals Gerontol. – Ser. A Biol. Sci. Med. Sci. 64, 807–815. doi:10.1093/gerona/glp027

Weir, J., 2005. Quantifying test-retest reliability using the intraclass correlation coefficient and the SEM. J. Strength Cond. Res. 19, 231–240. doi:10.1007/978-3-642-27872-3_5

